# The use of vocal coordination in male African elephant group departures: evidence of active leadership and consensus

**DOI:** 10.1101/2024.05.31.596833

**Authors:** Caitlin E. O’Connell-Rodwell, Jodie L. Berezin, Alessio Pignatelli, Timothy C. Rodwell

## Abstract

Group-living animals engage in coordinated vocalizations to depart from a location as a group, and often, to come to a consensus about the direction of movement. Here, we document for the first time, the use of coordinated vocalizations, the “let’s go” rumble, in wild male African elephant group departures from a waterhole. We recorded vocalizations and collected behavioral data as known individuals engaged in these vocal bouts during June-July field seasons in 2005, 2007, 2011, and 2017 at Mushara waterhole within Etosha National Park, Namibia. During departure events, we documented which individuals were involved in the calls, the signature structure of each individual’s calls, as well as the ordering of callers, the social status of the callers, and those who initiated departure. The “let’s go” rumble was previously described in tight-knit family groups to keep the family together during coordinated departures. Male elephants are described as living in loose social groups, making this finding particularly striking. We found that this vocal coordination occurs in groups of closely associated, highly bonded individuals and rarely occurs between looser associates. The three individuals most likely to initiate the “let’s go” rumble bouts were all highly socially integrated, and one of these individuals was also the most dominant overall. This suggests that more socially integrated individuals might be more likely to initiate, or lead, a close group of associates in the context of leaving the waterhole, just as a dominant female would do in a family group. The fact that many individuals were often involved in the vocal bouts, and that departure periods could be shorter, longer, or the same amount of time as pre-departure periods, all suggest that there is consensus with regard to the act of leaving, even though the event was triggered by a lead individual.

## Introduction

Group-living animals rely on vocalizations to identify and communicate with individuals at a distance, assess reproductive status, facilitate social interactions, and coordinate movement (Bousquet et al. 2011; O’Connell-Rodwell et al. 2012; Poole et al. 1988; Stewart & Harcourt 1994; Walker et al. 2017). Coordinating movement confers advantages, such as not getting separated from the rest of the group (Boinski & Campbell 1995; Walker et al. 2017), ensuring group members have met their physiological needs (e.g., food and water) (Sueur et al. 2010), and conserving energy by moving in relative synchrony, minimizing localization effort if separated (Black 1988; Boinski 1991). Mountain gorillas and redfronted lemurs have pre-departure vocalizations called “grunts” (Sperber et al. 2017; Stewart & Harcourt 1994) and white-faced capuchins make pre-departure “trills,” (Boinski & Campbell 1995), that cause the entire group to get ready and then move from an area. In wild dog packs, the incidence of sneezing increases prior to departure, acting as a quorum to confirm the group is ready to depart (Walker et al. 2017).

Vocalizations in elephants contain information about sex (Baotic & Stoeger 2017), age and body size (Stoeger & Baotic 2016), condition, and social and reproductive status (Poole et al. 1988; Soltis et al. 2005). Information encoded within calls makes it possible to identify individuals (McComb et al. 2003; Soltis et al. 2005; Stoeger & Baotic 2016; Wierucka et al. 2021) as well as be used to single out specific individuals in a noun-verb combinatory call, often made at a distance (Pardo et al. *in press*). Elephant vocalizations are also used to coordinate action within family groups, often initiated by either the matriarch or another dominant female within the family (O’Connell-Rodwell et al. 2012; Poole 2011; Poole et al. 1988).

In elephant family groups, matriarchs have been described as leaders (Lee & Moss 2012) because they make decisions for their family and act as knowledge repositories based on their experiences (Mutinda et al. 2011). Matriarchs assess predator threats to determine when to act (McComb et al. 2011), and make foraging decisions and initiate movement (Mutinda et al. 2011), such as when to leave the waterhole (O’Connell-Rodwell et al. 2012).

While male elephants are not considered group living animals, many individuals appear to spend a lot of time in all-male groups (Chiyo et al. 2011; Evans & Harris 2008; Goldenberg et al. 2014; Lee et al. 2011). However, little research has been conducted to assess the potential of male elephant coordination or active leadership. While male elephants have weaker associations within all-male groups than females do within their families (Archie et al. 2006; Chiyo et al. 2011), their social lives are very complex. Male elephants have been found to establish dominance hierarchies within social networks (O’Connell-Rodwell et al. 2011) and gather in large groups where males of all ages prefer to associate with older, mature males (Evans & Harris 2008). Preference for older males is likely attributed to older males taking on similar roles as matriarchs: older males aid in maintaining social cohesion (Chiyo et al. 2011), mediate aggressive behaviors (Allen et al. 2021; Slotow et al. 2000), and provide ecological information about resource location and effective navigation through the environment (Allen et al. 2020).

Individuals within bonded social groups coordinate their behavior and activities, which serves to maintain social stability through the use of physical interactions and vocalizations (Seltmann et al. 2013). Male elephants form social groups with older, more dominant males, sometimes appearing to take on a mentor or leadership role (Allen et al. 2020). While the evidence presented from photographs appears to support passive leadership, i.e. younger individuals following older individuals (Allen et al. 2020), we propose that some highly associated individuals, and especially the high-ranking male within an extended social network, may engage in active leadership tactics by initiating group departures vocally.

In this study, we document the use of “let’s go” rumble (LGR) vocalizations within bonded groups of male African elephants. We also show that these LGR events are mostly initiated by the most socially integrated individual. The initial LGR vocalization within a waterhole visit event triggers a series of highly synchronized and coordinated vocalizations within repeated bouts, a patterning that Poole (2011) refers to as cadence, as the dynamic resembles, and likely is a form of conversation to reach consensus. These bouts are often led by the most dominant individual and responded to by highly bonded individuals. We discuss the value of having such a vocal tool to trigger action and coordinate movement of a group of associates, as well as highlighting the evidence for, and implications of, active leadership of highly socially integrated individuals within male elephant groups.

## Materials & Methods

### Field site and elephant identification

Data were collected during June-July field seasons in 2005, 2007, 2011, and 2017 at Mushara waterhole (hereafter referred to as Mushara) in Etosha National Park, Namibia. Mushara is located within a 0.22 km^2^ clearing. Data were collected from an 8-meter-tall research tower, located approximately 80 meters from the waterhole. The waterhole is fed by a permanent, artisanal spring, and is the only stable source of water within 10 km^2^, making it an important resource during the dry season. For additional details about the field site, see recent publications (Berezin et al. 2023; O’Connell-Rodwell et al. 2022; O’Connell-Rodwell et al. 2022).

Elephants have been individually identified at Mushara since 2004 using unique, recognizable morphological characteristics such as ear tear patterns, tail hair configurations, tusk size and shape, and scarring. Elephants were assigned to age classes based on overall body size, shoulder height, hindfoot length, and skull and face morphometrics (Moss 1996; O’Connell-Rodwell et al. 2022).

Keystone individual (the most socially integrated and dominant individual in a population) identification using social network and dominance hierarchy analyses was described recently (O’Connell-Rodwell et al. 2024) and will be summarized in brief. For the social network analysis, we constructed association networks based on co-presence at the waterhole during field seasons. Weighted matrices of dyad-level association indices were built based on the Simple Ratio Index of association, ranging from 0-1, with higher indices representing individuals who are closely associated (Cairns & Schwager 1987; Whitehead 2008).

For the dominance hierarchy, we used dyad-level displacement (when an individual forces another to change his position; (O’Connell-Rodwell et al. 2011), to construct an ordinal hierarchy using the normalized David’s Score (David 1987; de Vries et al. 2006; Gammell et al. 2003).

David’s Score is calculated using the proportion of wins or losses across all dyads an individual is present in, while also considering the total number of dominance interactions observed. The highest values are associated with those who most consistently win contests. One individual (#22) had the highest average eigenvector centrality (most socially-integrated) and the highest dominance rank of all individuals included in the analysis across five years (2007 to 2011).

### Data acquisition

We recorded LGR vocalization events in the context of male elephants leaving Mushara waterhole. For each LGR event, we quantified the temporal spacing of the event, the onset of the departure period, the characteristics and individuality of LGR rumbles, the level of association between individuals that engaged in the bouts, and the behavior patterns within events, as well as bout initiation and serial participation of known individuals within the bouts.

Behavioral data and vocalization recordings were collected opportunistically during the evening and night (approximately 5:00 p.m. to 2:00 a.m.) when ambient sound and wind shear was low enough to record extremely low-frequency male vocalizations made in the range of 11 Hz. After dark, light-enhancing technology was attached to a standard HD video recorder and 3x magnification was used to visually identify individuals and document their behavior. In the new moon period, an infrared spotlight was also attached to the recorder to enhance visibility of tusks, ear tears and tail hair for individual identification.

Vocalizations were recorded using a Neumann Km131 microphone (Berlin, Germany) placed 20 meters from the waterhole, powered remotely via a 12-volt battery in the field tower.

Vocalization data collected in 2005-2011 was recorded using a TEAC DAT digital recorder, and in 2017, a Sound Devices solid-state digital recorder (Reedsburg, Wisconsin, USA) was used.

All vocalizations recorded were logged by date, time, type, and social context, including all individuals involved, the locations of callers, and those participating in the vocal bouts when known. Calls were flagged when it wasn’t possible to tell who the caller was, due to an obstruction (another elephant, the tower, or too far away to distinguish which individual was ear-flapping), or overlap with another caller, and were labeled as unknown.

Events were described as a period when a group of male elephants entered the clearing (from the forest) to the time when they departed the clearing. The criteria used to select events was as follows: 1) audio recordings were captured for the full event (from arrival to departure), 2) males arrived and departed together, and 3) females were not present during any time of the event, nor any other behaviorally impactful disturbances. Events were divided into pre-departure and departure periods following protocols described in O’Connell-Rodwell et al. (2012): pre-departure began when the elephants entered the clearing and was defined by greetings between males and drinking water, and ended when the departure period began. Departure began when a known male initiated the behavior associated with the “let’s go” rumble (and could be heard in almost all cases, due to the proximity of the microphone to the caller at the waterhole, as well as low-frequency sounds being more easily detectible after dark, given the low wind shear and quiet background) and ended when all elephants left the clearing. The microphone was monitored remotely using headphones plugged into the recorder in the tower.

Behaviorally, “let’s go” rumble bouts were identified when a known male stepped away from the waterhole, stood still and rumbled while flapping his ears, and positioned facing away from the waterhole (O’Connell-Rodwell et al. 2012; Poole et al. 1988). This first rumble marked the onset of the departure period.

After the initial rumble was emitted, the individual repeated the vocalization, while remaining stationary, or while walking away from the waterhole. This initial LGR call, or sequence of repeated calls, then triggered a bout of coordinated responses from the rest of the bonded group, a pattern that Poole (2011) refers to as cadence. Each caller within the bout was noted by ear-flapping behavior, while standing stationary or walking out to follow the initiator. These bouts were recorded until the group hit the edge of the clearing.

### Acoustic analysis

Rumbles were analyzed using Raven Pro 1.6 (Cornell Lab of Ornithology, New York, USA) with a Hann window size of 65536, a hop size of 32768, with 50% overlap. The window size is larger than previous publications (Stoeger & Baotic 2016; Wierucka et al. 2021) to precisely identify the fundamental frequency and harmonics. However, this extremely precise frequency resolution comes at the cost of a lower time resolution. “Let’s go” bouts have slightly overlapping rumbles or any calls made within 2 seconds were considered within a single bout. For non-overlapping rumbles, the full rumble was selected. For rumbles that do overlap, only the non-overlapping section is selected. For this study, only slightly overlapping bouts were considered as part of the LGR bout, or departure conversation. Individual rumbles were assumed to not be part of the “let’s go” rumble bout sequence. For bouts with more than three rumbles, only the first three rumbles of each bout were considered in the acoustic analysis.

A combination of parameters were used to identify individuals: 1) field notes, detailing the behavioral observation noting the time of “let’s go” rumble behaviors and the corresponding times on the audio recorder; and 2) the rule of non-consecutive rumble criteria (O’Connell-Rodwell et al. 2012), where it is assumed that the same elephant cannot rumble twice in a row per bout (but could be caller #1 and #3). Where it was difficult to behaviorally discern between two individuals, principal components visualizations of rumble characteristics were used to identify unique individuals, except for one case where we did not have individual rumbles for two callers in order to know which individual was which within the analysis (Table 2, #105/#69 (1) and #105/#69 (2)).

Following the methodology of Wierucka et al. (2021), we measured five key acoustic parameters: Frequency 5% (frequency that divides the rumble into two frequency intervals containing 5% and 95% of the energy), Frequency 95% (frequency that divides the rumble into two frequency intervals containing 95% and 5% of the energy), Bandwidth 90% (the difference between the 5% and 95% frequencies), Center frequency (divides the rumble into the two frequency intervals of equal energy), and Duration 90% (the differences between the 5% and 95% times) (abbreviated definitions reproduced from Charif et al. (2010) and Wierucka et al. (2021).

### Statistical analysis

To evaluate whether the onset of the “let’s go” rumble bouts trigger departure, we used a paired Wilcoxon Signed Rank test to assess whether the pre-departure time was significantly longer than the post-departure time, using the function “wilcox.test’ in the R stats package (R Core Team 2023). Similarly, a Wilcoxon Signed Rank test was also used to evaluate whether the number of rumbles significantly increased in the departure period (compared to the pre-departure period). Since longer events would be expected to have more rumbles, we calculated the rate of rumbles as the number of rumbles per minute in each period.

Next, we wanted to confirm that each “let’s go” rumble emitted contained a unique signature distinctive to each known individual, reproducing the methodology of Wierucka et al. (2021). Acoustic parameter data was normalized on a scale of 0 to 1 due to the different variable types, mean values of each variable, and disparate standard deviations. We used a Permutational Multivariate Analysis of Variance test (PERMANOVA) using the adonis function in the “vegan” package (Oksanen et al. 2022) with a Euclidean distance matrix of the frequency parameters. To confirm that differences were indeed due to the uniqueness of the calls between individuals, and not due to high within-individual variation, we tested for the homogeneity of variances using the betadisper function in the “vegan” package, followed by an ANOVA test.

Lastly, we assessed whether the males involved in “let’s go” events had significantly higher associations than those not involved in the “let’s go” events. Only association data from the 2007 field season was used, due to the large number of dyads observed, with data available for all individuals included in “let’s go” events. To increase the sample size of dyadic-relationships within “let’s go” events, we included five additional groups of individuals that were observed and acoustically recorded in a “let’s go” rumble event. These events could not be included in acoustic analysis, due to the lack of clear arrival times and audio recording of the entire event. Of the 26 individuals that came to the waterhole at least three times in 2007, there were a total of 223 unique dyads. Of these unique dyads, 64 dyads involved 17 highly associated individuals. 159 of these dyads involving 20 individuals, were not highly associated. Only the 17 highly associated individuals were involved in the “let’s go” rumble events. We used a Mann-Whitney U-test to assess for significant differences between the two groups of individuals (those observed in LGR events and those who are not), using the wilcox.test function in the “stats” package, with the alternative parameter set to “less.”

All statistical analyses and visualizations were performed using R statistical software (version 4.3.1) (R Core Team 2023), with significance set at an alpha level of α = 0.05.

## Results

### LGR Events and Temporal Spacing

The final acoustic analysis included data from 7 LGR events, with a total of 48 bouts and 122 analyzed rumbles (Table 1). A total of 19 individuals were recorded across the 7 LGR events (Table 2), with a mean group size was 4.9 individuals, with a range of 3 to 8 individuals (with only 7 individuals present across LGR events who did not vocalize). Nearly all the individuals involved in the LGR events were in the 3Q age class and older (25+ years old), with only 3 individuals in the 1Q (10-14 years old) and 2Q (15-24 years old) age classes.

**Table 1.** Description of “let’s go” rumble (LGR) events used in this study. Year, event number, number of bouts, number of rumbles per event, the duration of the event, number of males engaged in LGRs versus the total number within the group, as well as who initiated and the social context.

| LGR Event | Year | # Bouts | # Rumbles Recorded (analyzed) | Total Event Time (minutes) | # Elephants Vocalizing | # Elephants Present | Initiator of event | Keystone male present? |
| --- | --- | --- | --- | --- | --- | --- | --- | --- |
| 1 | 2005 | 8 | 22(20) | 55.0 | 4 | 4 | #22 | Yes |
| 2 | 2007 | 10 | 26(24) | 98.2 | 6 | 6 | #22 | Yes |
| 3 | 2007 | 1 | 3(3) | 50.0 | 3 | 5 | #67 | No |
| 4 | 2007 | 3 | 9(8) | 37.3 | 3 | 4 | #22 | Yes |
| 5 | 2011 | 5 | 17(13) | 31.9 | 3 | 3 | #84 | No |
| 6 | 2011 | 11 | 29(26) | 57.0 | 5 | 8 | #46 | No |
| 7 | 2017 | 10 | 32(28) | 56.0 | 4 | 5 | #46 | No |

**Table 2.** Individual known elephants, including the years that rumble vocalizations were recorded, along with a description of the number of rumbles recorded and rumble characteristics.

| Individual | Years recorded | Rumble count | Mean frequency 5% (Hz) | Mean fundamental frequency (Hz) | Mean center frequency (Hz) | Mean Frequency 95% (Hz) | Mean Bandwidth 90 (Hz) | Mean Duration (seconds) |
| --- | --- | --- | --- | --- | --- | --- | --- | --- |
| #18 | 2005, 2007, 2017 | 15 | 11.04 | 13.29 | 16.89 | 70.95 | 59.91 | 4.57 |
| #22 | 2005, 2007 | 23 | 12.93 | 13.99 | 28.80 | 78.02 | 65.09 | 4.04 |
| #25 | 2007 | 6 | 14.16 | 14.29 | 31.25 | 81.01 | 66.85 | 4.22 |
| #46 | 2007, 2011, 2017 | 23 | 10.89 | 12.99 | 19.52 | 82.22 | 71.33 | 4.55 |
| #48 | 2007 | 2 | 8.06 | 13.78 | 28.56 | 106.20 | 98.15 | 2.49 |
| #61/#132 (1) | 2017 | 8 | 9.52 | 13.07 | 13.73 | 45.04 | 35.52 | 3.94 |
| #61/#132 (2) | 2017 | 1 | 13.18 | 13.28 | 67.38 | 96.68 | 83.50 | 4.16 |
| #65 | 2007, 2011 | 7 | 11.72 | 13.09 | 20.30 | 64.45 | 52.73 | 4.25 |
| #67 | 2007 | 1 | 7.32 | 21.75 | 24.90 | 146.48 | 139.16 | 2.53 |
| #76 | 2011 | 7 | 13.18 | 13.64 | 25.78 | 77.01 | 63.83 | 5.96 |
| #83 | 2011 | 4 | 10.25 | 12.17 | 19.41 | 76.54 | 66.28 | 2.44 |
| #84 | 2011 | 5 | 11.43 | 12.49 | 24.90 | 87.31 | 75.88 | 4.32 |
| #105/#69 (1) | 2007 | 1 | 16.11 | 16.50 | 74.71 | 96.68 | 80.57 | 3.24 |
| #105/#69 (2) | 2007 | 6 | 9.52 | 13.87 | 23.68 | 87.40 | 77.88 | 3.01 |
| #118 | 2011 | 1 | 17.58 | 16.56 | 36.62 | 92.29 | 74.71 | 5.58 |
| #119 | 2011 | 5 | 11.13 | 13.63 | 20.22 | 85.55 | 74.41 | 3.89 |
| #146 | 2011 | 4 | 10.99 | 12.71 | 24.17 | 82.76 | 71.78 | 3.01 |
| Unk1 | 2005 | 2 | 10.99 | 11.48 | 17.58 | 24.90 | 13.92 | 2.94 |
| Unk2 | 2005 | 1 | 10.25 | 13.97 | 14.65 | 89.36 | 79.10 | 4.13 |

LGR events were defined by the pre-departure period, which was the arrival of a group of male elephants at the waterhole where they drank and socialized, followed by the departure period which was initiated by the onset of a “let’s go” rumble. Three rumble types were observed during these events, namely the first single call by the initiator (Fig. 1A) which triggered the highly synchronized and coordinated bouts that contained slightly overlapping rumbles emitted within bouts (Fig. C), by some or all of the individuals within the group at the waterhole.

**Figure 1.**
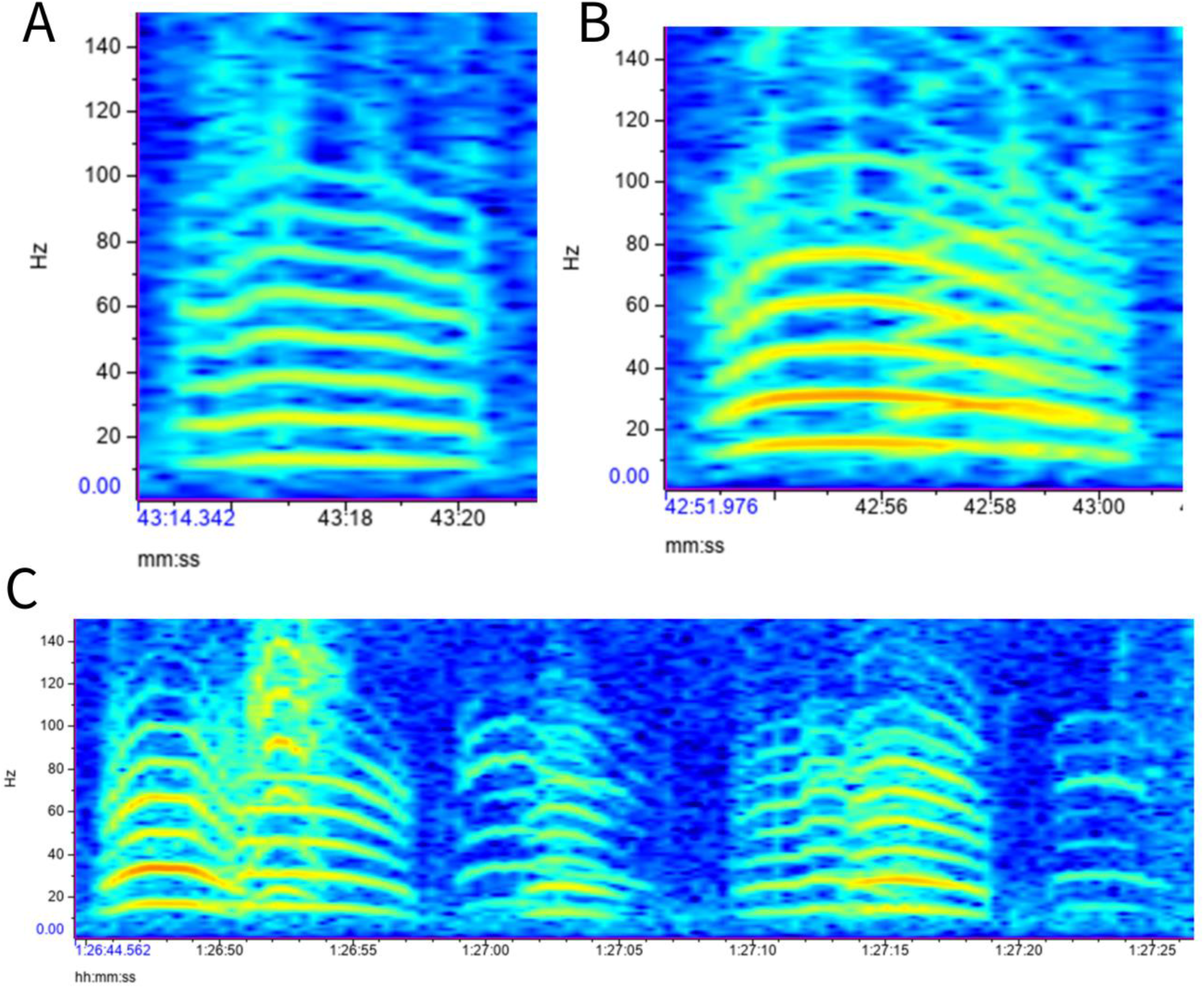
Three spectrograms depicting three different male elephant coordinated departure rumble vocalizations. (A) The “let’s go” rumble (LGR) is a single call from the initiator that triggers the LGR bouts (B) The LGR duet between the initiator and a close associate. (C) LGR bouts emitted by the rest of the group in response to the LGR trigger.

Sometimes, the initial vocalization was followed by an overlapping, “duet” call by the initiator and close associate (Fig. 1B). A spectrogram of an excerpt from event 2 depicts vocalizations in real time (Fig. 2). Sometimes, the initiator emitted a call, but did not get an immediate response, and proceeded to call several more times and even started walking away from the waterhole, before others responded (Fig. 2). In the example depicted in Figure 5, of a subset of rumbles, the keystone male, #22 emitted two LGR before triggering several bouts of rumbles that he almost always led (Fig. 5). The repeated vocal bouts resulted in the act of leaving the waterhole, most often as a group, though sometimes there were stragglers that return to the water for one more drink before following the rest of the group out of the clearing.

**Figure 2.**
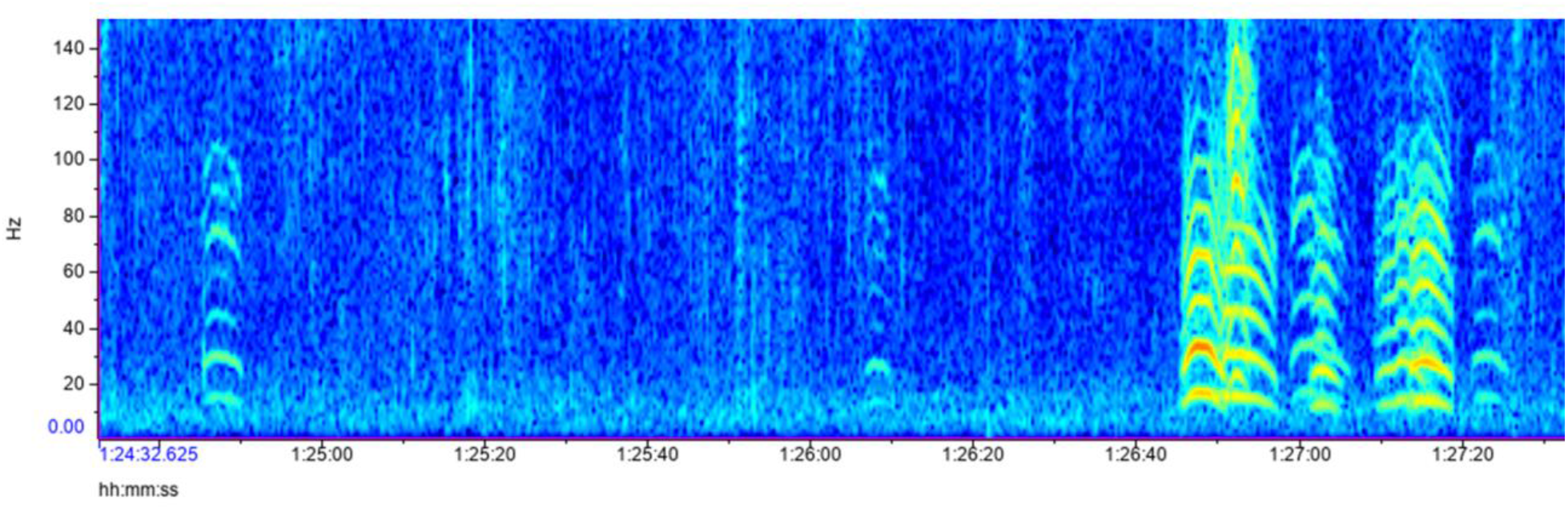
This spectrogram depicts a portion of the male elephant lets go rumble event within the post-departure period. The initiator emits a single LGR call, and the call sometimes gets repeated several times (in this case, the last two single calls in a series of repeated single calls are depicted, prior to triggering the LGR bout responses from the rest of the group). After emitting a LGR upon leaving the waterhole, the initiator did not get a response (at 1:24:90), the lower intensity dB level depicting that the initiator is heading away from the group and repeats the call again about a minute later (1:26:10). When he does not get a response the second time, he returns to the group at the waterhole and emits another LGR, this time triggering a series of LGR bouts emitted by the rest of the initiator’s close associates in response.

Four of the seven LGR events had a longer pre-departure than departure period (Fig. 3). The median pre-departure time (30.0 ± 9.68 minutes, range = 15.67, 42.50) was longer than the departure time (21.67 ± 16.5 minutes, range = 4.91, 55.97) but was not significant (Wilcoxon Signed Rank test *p* = 0.469, effect size *r* = 0.32, magnitude = moderate). Event 2 was unique in that there was an initial bout, then 43 minutes passed before a series of 9 bouts occurred in quick succession. During the 43 minutes between the first bout and the series, 14 individual rumbles were vocalized by the keystone individual (#22). When tested without event 2, the median times were still not significantly different (Wilcoxon Signed Rank test *p* = 0.313, effect size *r* = 0.47, magnitude = moderate).

**Figure 3.**
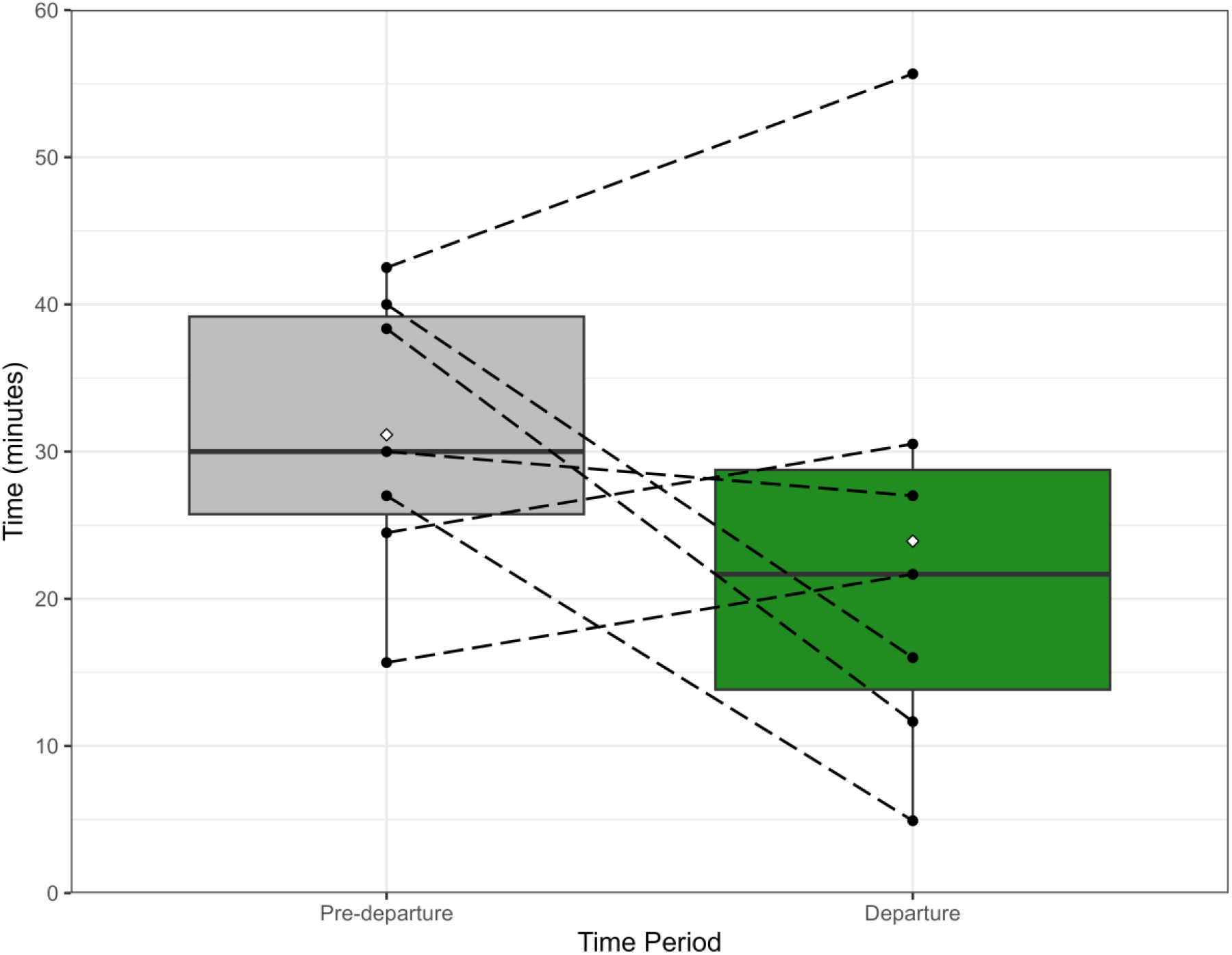
Total time (in minutes) the pre-departure and departure periods for the seven LGR events. Dashed lines connecting two solid black circles represent an event’s corresponding pre-departure and departure time, to visualize the relationship between the amount of time groups were present in each period. Thick horizontal bars represent the median, while the white diamond shape within each plot represents the mean. The vertical length of the box represents the interquartile range and the vertical lines are the minimum and maximum values. The median pre-departure time was longer than departure time, but the periods did not differ significantly (Wilcoxon Signed Rank test p = 0.469). For four events, the pre-departure time was longer than the departure time, while the other three events had longer departure periods.

The rate of rumbles in the departure period was significantly higher than the pre-departure period (Wilcoxon Signed Rank test *p* = 0.016, effect size *r* = 0.89, magnitude = large). For all events, pre-departure periods were silent, with no vocalizations recorded. The median rate of rumbles per minute in the departure period was 0.84 ± 1.12 (range = 0.26, 3.46; mean = 1.25).

Across all events, the mean (± SD) number of bouts per departure period was 6.86 ± 3.89 with a range of 1 to 11. The mean (± SD) number of rumbles was 19.71 ± 10.67 with a range of 3 to 32, while the mean (± SD) number of rumbles per bout was 2.88 ± 0.96 with a range of 2 to 6. The mean (± SD) duration of bouts was 10.54 ± 3.81 seconds with a range of 3.77 and 19.51 seconds. The average time between bouts was 156.55 ± 405.40 seconds (2.61 ± 6.76 minutes) with a range of 2.80 and 3624.23 seconds (0.047 to 43.73 minutes).

### Rumble characteristics and individual differences

The mean duration of rumbles was 4.15 seconds (SD = 1.42) and the mean Frequency 5% was 11.53 Hz (SD = 2.31). Additional rumble characteristics are presented in Table 3. We found significant individual differences in the five acoustic parameters for the 19 individuals included in the study (PERMANOVA *R*^2^ = 0.522, p = 0.001; Table 3). Further, the homogeneity of variance assumption was not significant, (*F* = 1.34, DF = 18, *p* = 0.206), indicating that the differences between individuals were not due to large within-individual variation but due to inter-individual variation.

**Table 3.** Rumble parameter characteristics. SD = standard deviation; Min = minimum frequency; Max = maximum frequency.

| Parameter | Mean | Median | SD | Min | Max |
| --- | --- | --- | --- | --- | --- |
| Duration (seconds) | 4.15 | 4.13 | 1.42 | 1.51 | 8.89 |
| Bandwidth (Hz) | 65.49 | 67.38 | 17.74 | 11.72 | 139.16 |
| Center Frequency (Hz) | 23.26 | 24.17 | 9.01 | 11.72 | 74.71 |
| Frequency 5% (Hz) | 11.53 | 11.72 | 2.31 | 5.86 | 17.58 |
| Frequency 95% (Hz) | 77.016 | 79.10 | 17.65 | 21.97 | 146.48 |

### Associations, dominance, and the keystone individual

Of all the frequent visitors to Mushara in 2007, individuals within LGR groups had a mix of association levels amongst its members, where some individuals had high association strengths, and others had low. Males involved in LGR events (highlighted in yellow; Fig. 4) had significantly higher association indices than those that did not engage in LGR events (highlighted in blue; Fig. 4) (Mann-Whitney U test *p* = 0.0001, median difference = 0.05, effect size = 0.26, magnitude = small). The median index for those involved in an LGR event was 0.16 ± 0.17 (mean = 0.21, range = 0.04 to 0.92), while the median for those not observed in an LGR group was 0.11 ± 0.07 (mean = 0.12, range = 0.04 to 0.36).

**Figure 4.**
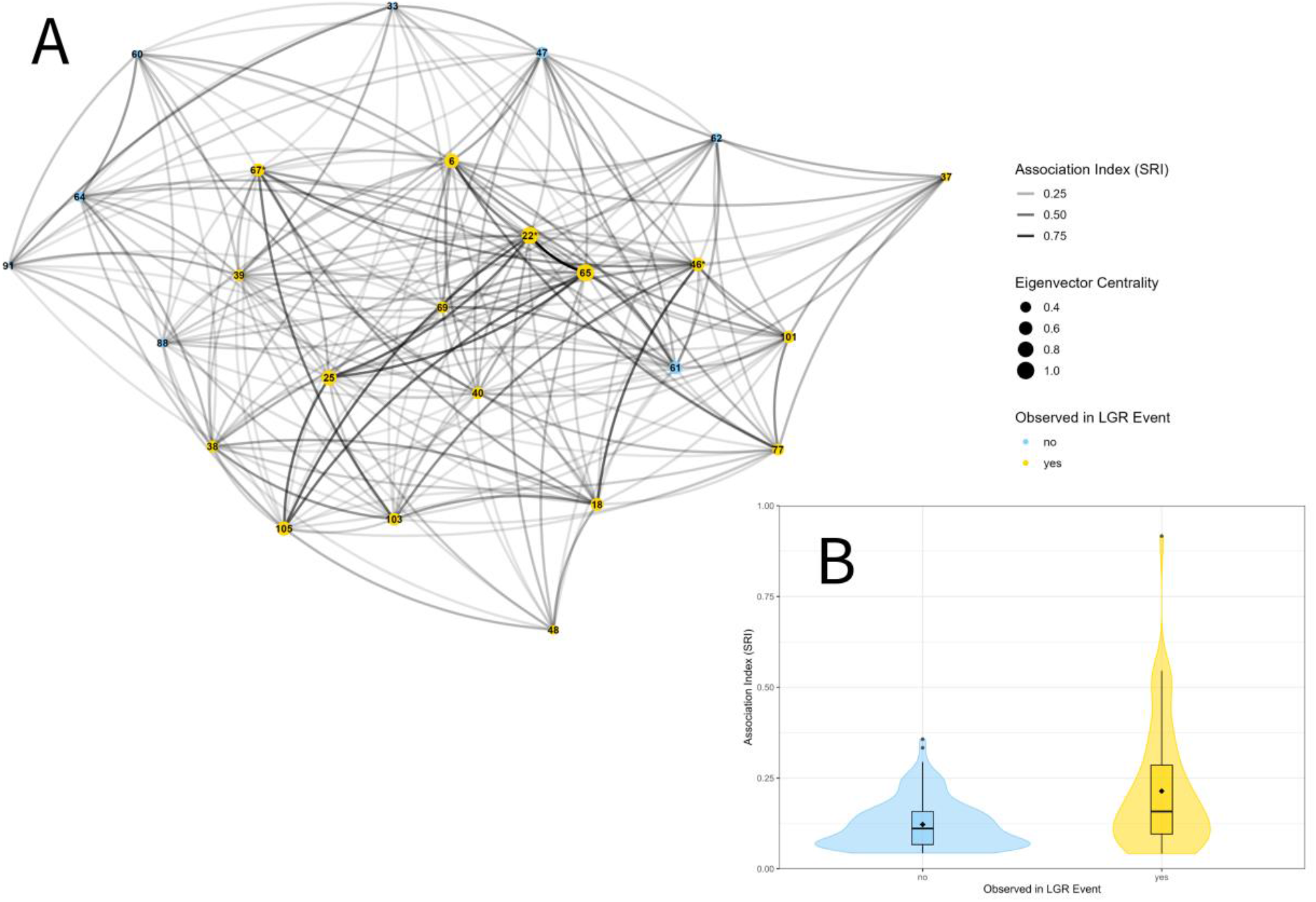
Association indices for individuals involved in LGR events (yellow) and those who are not (blue). (A) Social network for 2007 for n = 26 individuals. The lines between individuals represent the strength of association, binned into four levels, ranging from 0 to 1. Circles represent individuals and the size of the circle represents their centrality within the social network, binned into four levels. Three of the LGR event initiators are marked with asterisks next to their ID numbers. (B) Violin plot of the association indices for individuals not observed in an LGR group (n = 159 dyads) and those who were observed in an LGR group (n = 64 dyads). The shape of the violin plot is a visual representation of the distribution of the data. The box plots provide further details about the data distribution where the thick black line represents the median and the black diamond represents the mean. The length of the box is the interquartile range, with the vertical lines representing the minimum and maximum values, and filled circles represent outliers.

**Figure 5.**
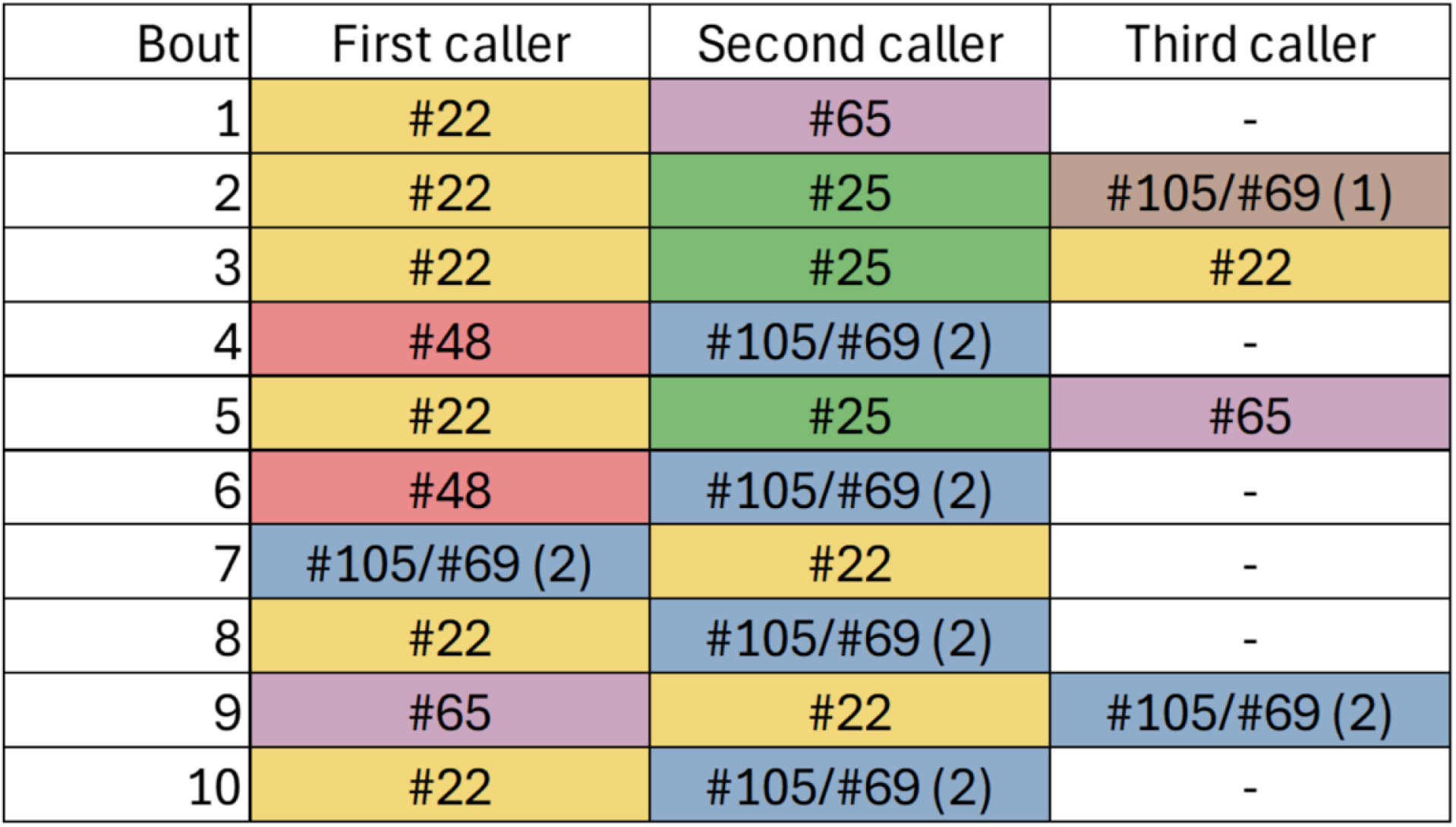
Example of the order of callers (rumbles) in each bout for event 2. The keystone male (#22) initiates most of the bouts in this event, while the other initiators were all mid-ranking (Fig. 6).

For the three events where he was present, the keystone male (#22) initiated the departure of the group by emitting a LGR 61.9% (13/21) of the time and was always the first caller in the LGR bouts. When he was present, six (of nine) other individuals in his groups are also initiators, but they only initiated bouts 1 or 2 times each, making the keystone male 1.6 times more likely to initiate than any other individual, when he was present. Across all events (when #22 was present and when he was not), 12 of the 19 individuals initiate bouts. When the keystone male was not present, one individual (#46) initiated 54.5% (12/22) of the bouts in the three events he was present in. All other individuals initiated five times or fewer (for example, see Fig. 5).

Males #22, #46, and #67 had high centrality rankings of 1, 10, and 14, respectively, out of 49 individuals evaluated (data was not available for male #84, the initiator of event 5). Of these three individuals, only male #22 was the highest ranked in the dominance hierarchy overall, while males #46 and #67 were mid-ranking overall and not the highest ranked members in their respective LGR groups (Fig. 6).

**Figure 6.**
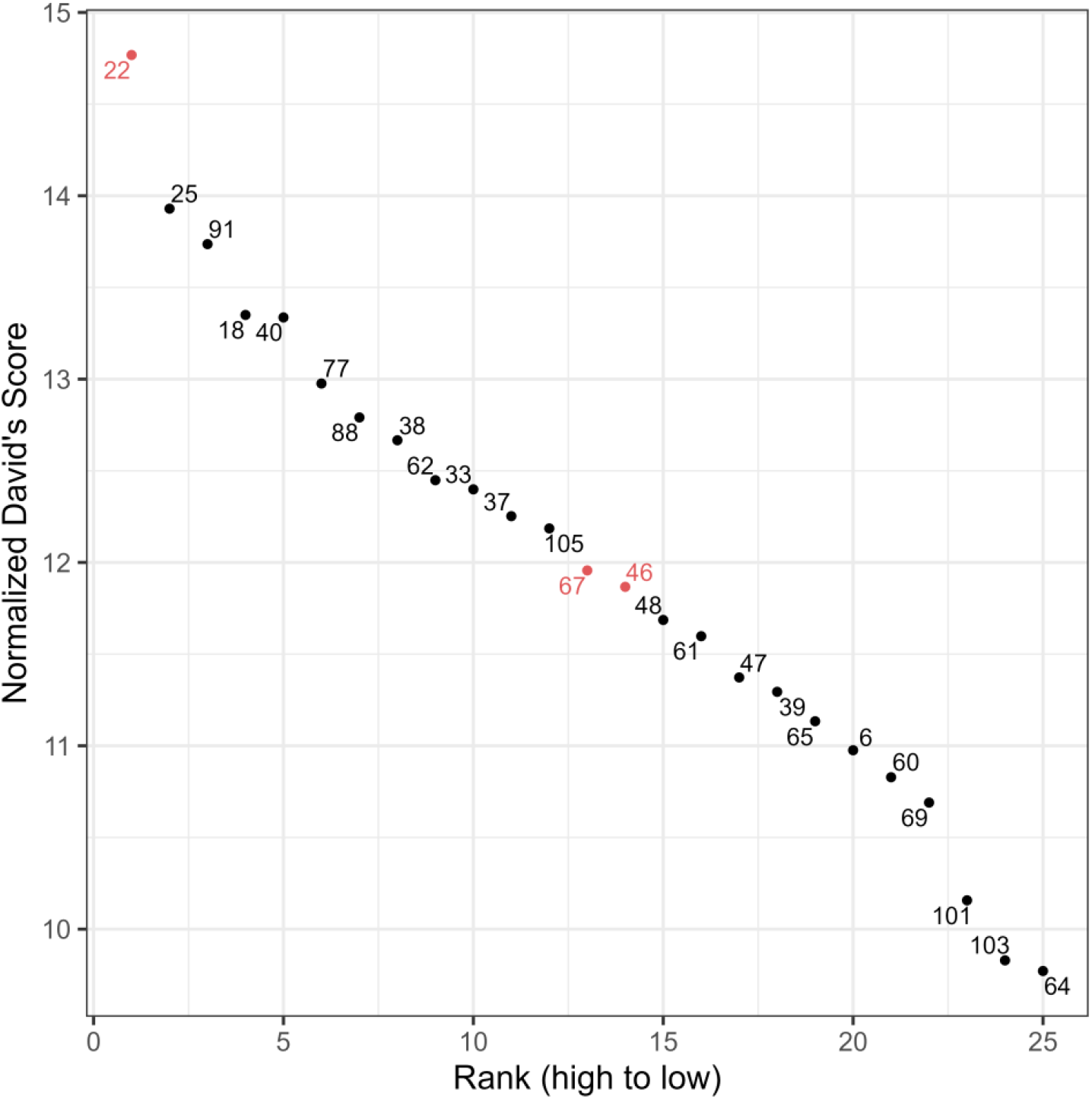
Dominance hierarchy for 2007, highlighting the three initiators for which data was available (in red). A total of 25 non-musth individuals were included in the analysis. Male #22 (the keystone individual) was ranked 1, while males #67 and #46 were ranked 13 and 14, respectively.

## Discussion

Since male elephants have been described as living in loose groups of associates (Archie et al. 2011; Chiyo et al. 2011), it is surprising to document them engaging in highly coordinated vocal behavior, used to coordinate departures from the waterhole as a group of associates, just as group-living animals do. And even more surprising, is that they do so with vocal patterning and synchrony (Fig. 1 and Fig. 2) previously only described in females living within family groups (O’Connell-Rodwell et al. 2012; Poole et al. 1988) as part of a departure conversation, or cadence (Poole 2011). To add to these surprising findings is the fact that this vocal coordination during departure only occurs within male groups that have strong associations and are much rarer between loose associates (Fig. 4).

This solicitous behavior suggests much deeper relationships than random meet ups at a waterhole while drinking, whereby individuals might engage in social interactions with bonded associates, and from there, perhaps passively follow a dominant or socially integrated individual upon departure. This vocal coordination among associates was also found in bonobos, whereby more bonded individuals were more effective at coordinating group action (Levrero et al. 2019), and adult male Barbary macaques most frequently recruited those who they had affiliative relationships (Seltmann et al. 2013). Although the level of dyadic associations varied in some male elephant groups—some individuals having low associations—each individual had a stronger association with at least one other individual in the group. Lending further evidence to the idea that these vocal bouts, or conversations, expedited departure is the fact that bonded groups that engaged in LGR bouts had more coordinated departures than loose affiliates.

The most intriguing aspect of these findings is that three of the “let’s go” event initiators (males #22, #46, and #67) were highly socially integrated (central) within the association network (Fig. 4), and only one of those individuals was also highly dominant overall (male #22; Fig. 6), all three being nearly fully mature (> 25 years old; #67) or fully mature adults (> 35 years old; #22 and #46)(O’Connell-Rodwell et al. 2022). Species social structure is thought to impact the coordination of movement (Seltmann et al. 2013), but results have been inconclusive as to who has the most social influence (Petit & Bon 2010). For example, social integration and maturity were important for coordinated movement in cattle (Šárová et al. 2013; Sueur et al. 2018). Being an adult, high-ranking male was important for Barbary macaques (Seltmann et al. 2013). And, lastly, dominance rank was the most important for successive rallying and departure for African wild dogs (Walker et al. 2017).

For male African elephants, our results suggest that dominance might not be the most important quality for male elephants in the coordination of departure, but rather social integration, maturity, and bondedness. Socially integrated individuals are thought to act as sources of social information (King & Sueur 2011), due to the quantity of connections within their network.

Central individuals might also have greater access to information (Palacios-Romo et al. 2019), making them more attractive as companions than less socially integrated individuals. For example, in male elephants, dominance hierarchies are constructed based on displacements at the waterhole, thus, being a dominant male often does not necessarily convey to others that an individual has knowledge about the social or physical environment.

Socially integrated individuals were the most likely to initiate the LGR events, but several other individuals initiated individual bouts within the events (Table 1). Additionally, a majority of the individuals in the group participated in the bouts (Table 1), suggesting that the final decision of when to depart is shared in a consensus (Sueur & Petit 2008). Collective decision-making is thought to be more accurate than a decision made with a lack of consensus, since it’s based on the knowledge of many individuals (Conradt & Roper 2005). For our male groups, the individuals who participated in the vocal bouts were all at least 25 years old (3Q age class; with the exception of individual #65; Table 2), all of whom would have decades of shared knowledge. Further, even the individuals who did not participate in the vocalizations (many of whom were mature adults) are considered to be part of the decision-making process just by following and “agreeing” non-vocally to the decision being made by the other individuals in the group (Conradt & Roper 2005).

Interestingly, the departure period was not significantly different from the pre-departure period, and three of the seven events had longer departure times than pre-departure (Fig. 3). In contrast to family groups where the matriarch has the most knowledge of the environment (McComb et al. 2001; McComb et al. 2011; Mutinda et al. 2011), the adult male elephants in our LGR groups likely all have similar repositories of environmental knowledge and are independent adults. As such, the initiators of the LGR events likely have less “control” than a matriarch might have over her family group and might require the males to have longer periods of decision-making, contributing to our observed longer departure periods. Future research might focus on the degree to which group size, rumble rate, or level of bondedness might impact departure duration.

We found a significant increase in the rate of rumbles and coordinated bouts in the post-departure versus the pre-departure period, where all events had zero rumbles in the pre-departure period. These results contrast with previous findings in female elephants where there were considerably more vocalizations made in the pre-departure period (O’Connell-Rodwell et al. 2012) than we observed in the male groups. Male elephants are described as being less vocal overall than females (reviewed in Morris-Drake & Mumby (2017)), which likely explains why there were so fewer vocalizations in the pre-departure period. Since there are many more individuals to have to rally, it makes sense that the females are more vocal in reaching consensus from other dominant females and their core families.

These results offer the first evidence of active leadership in male African elephants, whereby socially integrated and/or dominant individuals, actively determine the departure time and direction for the group, just as matriarchs do. A leader, or active leader, is defined as one who solicits those to follow them and exerts social influence over a group by means of their dominance rank, social position, experience, or a specific behavior (King et al. 2009; Pyritz et al. 2011). In contrast, passive leadership occurs when an individual might be unintentionally leading (King et al. 2009; Pyritz et al. 2011), such as what was previously described in male elephants where younger individuals followed mature males (Allen et al. 2020).

This coordination among males within highly associated groups begs the question of what advantage individuals might have in maintaining a group’s integrity over time and space. Maintaining bonds within groups strengthens group cohesion (de Waal 1986), which for social males, could facilitate coalition behavior, thus providing a competitive advantage over resources, such as scarce waterpoints in an arid environment. This competitive edge over adversaries might outweigh having to share resources with associates (Conradt & Roper 2000) and also reduces competition over scarce waterpoints (O’Connell-Rodwell et al. 2011). Finally, this behavior might benefit genetically related individuals involved in coordinated vocal departures, whereby shared social and environmental knowledge could serve to enhance reproductive benefits.

Further relatedness studies on associates may shed light on this possibility, but how individual males might discriminate paternity is an open question.

Finally, we found significant differences in rumble characteristics amongst individuals, supporting previous findings using similar methodologies (Stoeger & Baotic 2016; Wierucka et al. 2021). Our frequency 5% was extremely similar to Wierucka et al. (2021) and also fit within the range of the fundamental frequency previously reported (Baotic & Stoeger 2017; Poole et al. 1988; Stoeger & Baotic 2016). Further, our center frequency, duration, bandwidth, and frequency 95% fall within the range of those of Wierucka et al. (2021). These quantifiable differences in call structure between individuals is likely distinguishable by others within the cohort and could be used to keep track of who is calling at what distances, and monitor any adjustments in direction, while leaving the area to facilitate coordination.

It is also likely that the ‘let’s go” rumble differs acoustically from other vocalizations that male elephants produce, such as the musth rumble (Poole 2011; Poole et al. 1988), which tends to be a longer repeated call that does not elicit a response like LGR calls. Also LGR calls appear to have more modulation than the contact calls described by (Poole 2011; Poole et al. 1988; Wierucka et al. 2021). The antiphony of the LGR bouts, or cadence, as part of a decision-making process between bonded individuals is also very distinct, warranting further research into the “language” of male elephants.

## Conclusions

This study reports the first evidence of the use of vocal coordination in the departures of closely associated, male African elephants. We also provide the first evidence of active leadership in male elephants, whereby socially integrated individuals begin the departure period by actively recruiting their associate’s company during departure, using a “let’s go” vocalization. Most of the other group members participate in the decision making process, as far as the time and possibly the direction of the departure, similar to the negotiation of family groups (O’Connell-Rodwell et al. 2012), contrasting previous findings of passive leadership in males, where older males appeared to be unintentionally leading subordinates to resources (Allen et al. 2020).

These findings provide further support that mature males, and perhaps certain individuals such as those leading the LGR events here, are important for male elephant society (Allen et al. 2020; Allen et al. 2021; Chiyo et al. 2011; Goldenberg et al. 2014; Lee et al. 2011; Slotow et al. 2000).

Further studies are needed to understand the underlying advantages of such surprisingly coordinated vocal bouts within groups of male African elephants, the level of coordination and vocal manipulation, as well as conditions that evoke such behavior that has not yet been documented in other populations.

## Acknowledgements

The authors thank the Namibian Ministry of Environment and Etosha Ecological Institute for their support of this research. We also thank the contributing volunteers of Utopia Scientific and the Elephant Sanctuary for making the field work and data analysis possible. We are also grateful to Jason D. Wood for early help with acoustic analysis and training with Raven Pro.

